# A paradox of parasite resistance: Disease-driven trophic cascades increase the cost of resistance, selecting for lower resistance with parasites than without them

**DOI:** 10.1101/2021.11.05.466696

**Authors:** Jason C. Walsman, Alexander T. Strauss, Jessica L. Hite, Marta S. Shocket, Spencer R. Hall

## Abstract

Most evolutionary theory predicts that, during epidemics, hosts will evolve higher resistance to parasites that kill them. Here, we provide an alternative to that typical expectation, with an explanation centered on resource feedbacks. When resistance is costly, hosts evolve decreasing resistance without parasites, as expected. But with parasites, hosts can evolve *lower* resistance than they would in the absence of parasites. This outcome arises in an eco-evolutionary model when four conditions are met: first, resistance has a fecundity cost (here, via decreased foraging/exposure rate); second, resources increase during epidemics via trophic cascades; third, increased resources magnify the benefit of maintaining a fast foraging rate, thereby magnifying the cost of evolving a slower foraging/exposure rate (i.e., resistance); fourth, that amplification of the cost outweighs the benefit of resistance. When these conditions are met, hosts evolve *lower* resistance than without parasites. This phenomenon was previously observed in a mesocosm experiment with fungal parasites, zooplankton hosts, and algal resources. Re-analyzing this experiment produced evidence for our model’s mechanism. Thus, both model and experiment indicate that, via resource feedbacks, parasites can counterintuitively select against resistance.

## Introduction

Emerging infectious diseases threaten many host populations (Daszak et al. 2000; Vredenburg et al. 2010). Infectious diseases depress host density because they harm host fitness, often by increasing host mortality. Thus, infectious diseases can trigger conservation concerns (Roelke-Parker et al. 1996; Cooper et al. 2009), economic damage (Fry and Goodwin 1997), and human health crises (Munster et al. 2020). However, epidemics may depress host populations less when hosts evolve resistance (Altizer et al. 2003; Penczykowski et al. 2011). Resistance can evolve, for example, when host genotypes vary in their anti-infection resistance (inverse of transmission rate, ‘resistance’ hereafter). Such variation arises from mechanisms including differential immune investment (Valtonen et al. 2010), variation in exposure rates to parasites (Hall et al. 2010) or matching alleles between host and parasite (Agrawal and Lively 2002). More resistant genotypes become infected less often, and therefore suffer the effects of infection less often; all else equal, more resistant genotypes have higher average fitness. This benefit selects for resistance; as a result, average host resistance increases in the presence of parasites. Evolution of higher resistance then decreases the proportion of hosts infected and increases host density (Duffy and Sivars-Becker 2007).

Hosts can, instead, evolve *lower* resistance if resistance is costly (e.g., Duncan et al. 2011; Duffy et al. 2012). Resistant genotypes often pay a fitness cost in some other trait, e.g., fecundity (Fuxa and Richter 1998; Webster and Woolhouse 1999) or related resource use traits like foraging rate (Hall et al. 2010; Auld et al. 2013; Hall et al. 2012; Kraaijeveld and Godfray 1997). In the absence of parasites, resistance provides no benefit. Hence, any fecundity cost of resistance should select for lower resistance (Duncan et al. 2011). With parasites, selection then depends on the balance of the benefit and cost of resistance. The benefit of resistance depends on two factors. First, parasite abundance (related to prevalence) determines how often resistant host genotypes prevent infection compared to less resistant genotypes. Second, the mortality induced by infection (‘virulence’ hereafter) determines the benefit of each infection prevented. If the fecundity cost of resistance outweighs this survival benefit, selection favors lower resistance. Such evolution has occurred during some small epidemics in nature (Duffy et al. 2012). Regardless, according to conventional theory, adding disease should always increase the survival benefit of resistance (compared to no disease where there is no benefit). Thus, intuitively, epidemics are expected to select for some degree of resistance, slowing or reversing the direction of evolutionary change compared to parasite-free conditions. However, experimental epidemics in zooplankton (Fig. 1a) have exhibited evolution of *lower* resistance in host populations during epidemics (see Fig. 1b; Strauss et al. 2017). Such a result challenges this intuition, highlighting a need for novel mechanistic theory to explain this paradoxical phenomenon.

**Figure 1.**
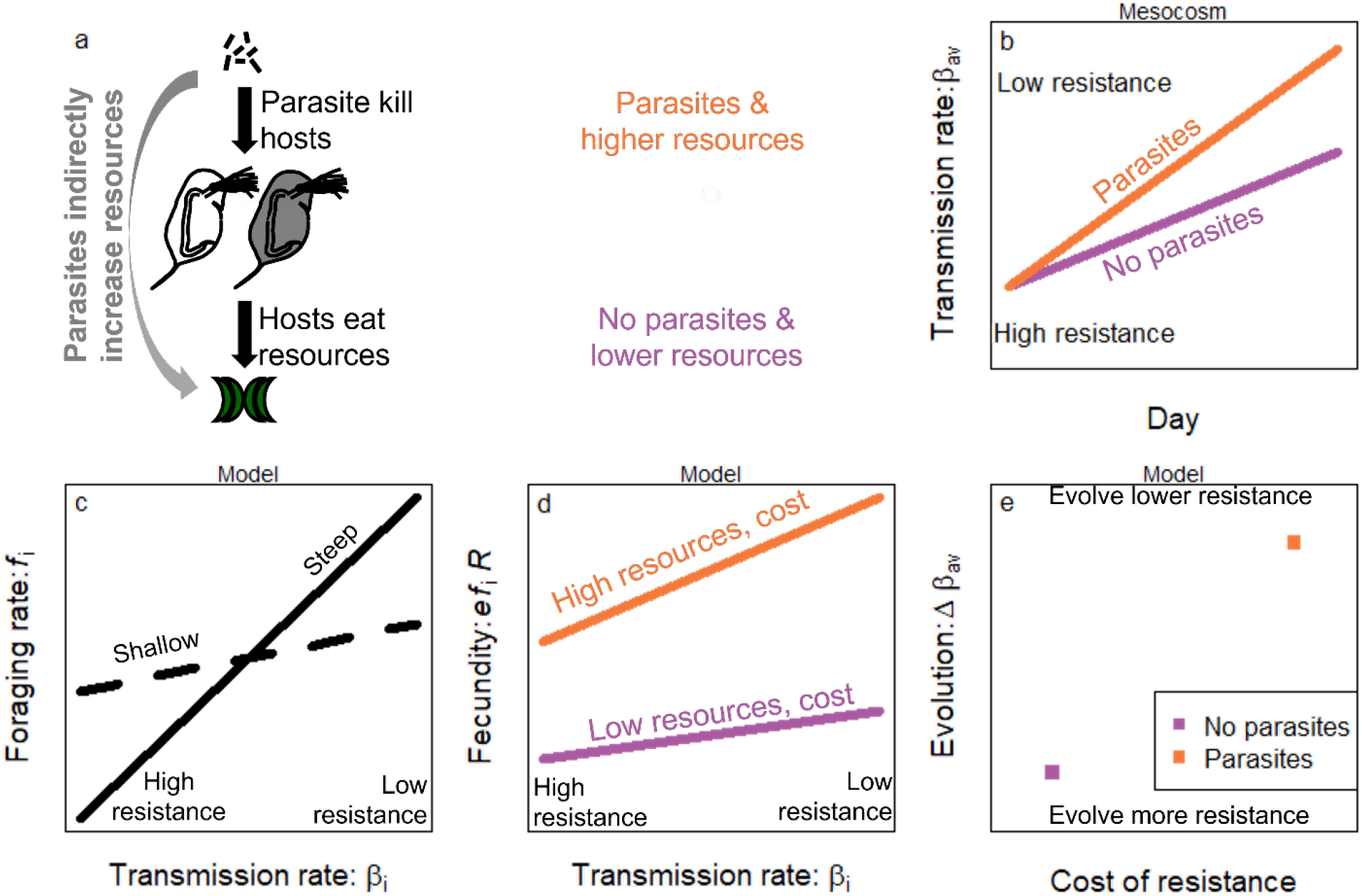
Conceptual overview. (a) Our model and the mesocosm both contain a simple community in which parasites kill hosts, which eat resources; thus, parasites indirectly increase resources in a trophic cascade. Orange denotes parasite presence and higher resources while purple denotes parasite absence and lower resources. (b) In the mesocosm experiment, host populations were observed evolving higher, average transmission rate (*β*_av_, lower resistance) with parasites (orange) than in their absence (purple). (c) Evolution of higher transmission rate can arise with positive covariance between *β*_i_ and foraging rate (*f*_i_). Genotypes which benefit from higher resistance pay a fecundity cost with lower *f*_i_. This tradeoff may be steep (solid, more costly resistance) or shallow (dashed less costly resistance; panels d and e use the steep tradeoff). (d) The *β*_i_-*f*_i_ tradeoff creates a fecundity cost of resistance (line steepness) that increases with resource density, *R*. (e) The model predicts that parasites can increase the cost of resistance, explaining evolution of higher *β*_av_ during epidemics than without. Panels c-e have matching empirical data showing the same qualitative pattern (see below).

Conventional theory and the prediction of increased resistance during epidemics rests on an assumption of a constant cost of resistance. Yet in reality, ecological factors, such as the density of resources for hosts, can alter the cost of resistance (Boots 2011; Zeller and Koella 2017). In many cases, the fecundity cost of resistance decreases when resources increase (Boots 2011; Duncan et al. 2011) as high resistance, low fecundity genotypes benefit more from increased resources. But if low resistance, high fecundity genotypes benefit more from increased resource quantity or quality, the cost of resistance can increase with resources (Vale et al. 2015; Hall et al. 2012). More resistant bacterial genotypes exhibited a cost in terms of growth rate at low bacterial density (corresponding to high resources) but not at high bacterial density (low resources; Vale et al. 2015). Similarly, increased resource quality for zooplankton hosts increased the cost of resistance (Hall et al. 2012).

We focus on increased quantity of resources, which can increase the cost of resistance in our focal, zooplankton system due to link between foraging and parasite exposure. Many host species become infected while foraging (Philpott et al. 2004; Hall et al. 2007; Coulson et al. 2018). Such connections with exposure can create a genetic tradeoff linking foraging rate to transmission rate (Fig. 1c; Hall et al. 2010; Hall et al. 2012). Because foraging rate is linked to fecundity, this creates a fecundity-transmission rate tradeoff (Hall et al. 2010; Auld et al. 2013). The fecundity cost of resistance among genotypes is given by the slope of the relationship (Fig. 1d); e.g., with a steep slope, more resistant genotypes have much lower fecundity than less resistant genotypes. Given this foraging-rate based tradeoff, increased resources can multiplicatively increase the cost of resistance (higher slope of orange line than purple in Fig. 1d).

The effect of resources on resistance evolution matters when parasites drive trophic cascades during epidemics. Trophic cascades arise in a diversity of parasite-host-resource systems (Buck and Ripple 2017): when parasites kill hosts, they indirectly increase resource density. This resource increase can amplify the fecundity cost of resistance, potentially outweighing the mortality benefit of resistance. If true, this mechanism would explain how hosts could evolve decreasing resistance *faster* during epidemics than in their absence (Strauss et al. 2017). To evaluate this hypothesis, we model an obligately killing parasite and a resource that both respond to changes in the density of the host population. Host populations are composed of clonal genotypes; those genotypes have positive covariance between transmission rate and foraging rate. That covariance influences evolution by clonal competition during epidemics. Hence, evolution and ecology share the same time scale. In this eco-evolutionary model, two outcomes arise. If the foraging-resistance tradeoff has a shallow slope (dashed line in Fig.1c, i.e., resistance is less costly), hosts evolve resistance. With a steeper tradeoff, a strong trophic cascade drives evolution of lower resistance, potentially even faster with disease than without (Fig.1e). We also evaluate how nutrient supply and resource competitors can alter these outcomes by shaping how resources respond to disease.

We test these model predictions by re-analyzing the dynamics of fungal parasite, zooplankton host, and algal resource populations during experiment epidemics (Strauss et al. 2017). We complemented the experiment with trait measurements on individual hosts; we measured foraging rate of host genotypes at two resource levels to show how elevated resources would increase the fecundity advantage of fast foragers (i.e., fecundity cost of resistance). This critical piece of the mechanism aids interpretation of the mesocosm experiment. Clonal assemblages were chosen for the mesocosm so that host populations had steeper or shallower tradeoffs. The experiment also varied supply of nutrients (low nutrient results first presented here) and presence of a smaller zooplankton that competes for algae (*Ceriodaphnia spp*.). Outcomes of these other treatments broadly reinforces our mechanistic explanation of the importance of resource availability in driving the evolution of resistance during epidemics. In support of the model (and conventional theory), epidemics caused selection for resistance despite elevated resources in populations with the shallower (weaker) tradeoff. But for host populations with a steeper tradeoff (more costly resistance), large trophic cascades resulted in stronger selection against resistance with disease than without. In other words, hosts evolved to become even more susceptible to infection, as epidemics proceeded. Low nutrients and competitor presence may have minimized the effect of parasites on resources, as predicted, but not significantly. Together, model and experiment reveal how parasites can drive cascades that may be altered by nutrients and competitors; the resulting increased resources can drive hosts to evolve lower resistance to a deadly parasite than they would in disease-free conditions.

## Eco-evolutionary Modelling

### Eco-evolutionary Model

An eco-evolutionary model captures how parasites alter selection on resistance directly by killing hosts and indirectly through their interaction with resources. The eco-evolutionary model of the density of resources (*R*), susceptible hosts of clonal genotype *i* (*S*_*i*_), infected hosts (*I*_*i*_), free-living parasite propagules (*Z*), and a competitor species (*C*) is (see also Table 1):

**Table 1.** List of symbols for state variables (top) and traits and other parameters (bottom) in eco-evolutionary model (eq. 1). Default values are accompanied by ranges.

| Symbol | Meaning | Units | Value |
| --- | --- | --- | --- |
| $t$ | Time | day | 0-125 |
| $R$ | Density of resources | $\mu\text{g chl } a \cdot \text{L}^{-1}$ | Varies |
| $S_i$ | Density of susceptible hosts, $i$ | $\text{hosts} \cdot \text{L}^{-1}$ | Varies |
| $I_i$ | Density of infected hosts, $i$ | $\text{hosts} \cdot \text{L}^{-1}$ | Varies |
| $Z$ | Density of parasite propagules | $\text{parasites} \cdot \text{L}^{-1}$ | Varies |
| $C$ | Density of competitors | $\text{competitors} \cdot \text{L}^{-1}$ | Varies |
| $H_i$ | Total host density, $I_i + S_i$ | $\text{hosts} \cdot \text{L}^{-1}$ | Varies |
| $p_i$ | Prevalence of infection, $i$, $I_i / (I_i + S_i)$ | unitless | 0-1 |
| $S_T$ | Total susceptible host density, $\sum_{i=1}^n S_i$ | $\text{hosts} \cdot \text{L}^{-1}$ | Varies |
| $I_T$ | Total infected host density, $\sum_{i=1}^n I_i$ | $\text{hosts} \cdot \text{L}^{-1}$ | Varies |
| $H_T$ | Total host density, $I_T + S_T$ | $\text{hosts} \cdot \text{L}^{-1}$ | Varies |
| $p_{av}$ | Average prevalence, $I_T / (I_T + S_T)$ | unitless | 0-1 |
| $h_i$ | Frequency of genotype $i$, $\sum_{i=1}^n H_i / H_T$ | $\text{hosts} \cdot \text{L}^{-1}$ | 0-1 |
| $d$ | Background mortality rate, focal hosts | $\text{day}^{-1}$ | 0.05<sup>a</sup> |
| $d_C$ | Background mortality rate, competitor | $\text{day}^{-1}$ | 0.05 |
| $e$ | Conversion efficiency, focal hosts | $\text{hosts} \cdot \mu\text{g chl } a^{-1}$ | 0.14<sup>a</sup> |
| $e_C$ | Conversion efficiency, competitor | $\text{competitors} \cdot \mu\text{g chl } a^{-1}$ | 1.68<sup>b</sup> |
| $f_i$ | Foraging rate, focal hosts $i$ | $\text{L} \cdot \text{host}^{-1} \cdot \text{day}^{-1}$ | 0.024-0.044<sup>a</sup> |
| $f_{av}$ | Average foraging rate, $\sum_{i=1}^n h_i f_i$ | $\text{L} \cdot \text{host}^{-1} \cdot \text{day}^{-1}$ | 0.024-0.044<sup>a</sup> |
| $f_c$ | Foraging rate, competitor | $L \cdot \text{competitor}^{-1} \cdot \text{day}^{-1}$ | 0.0038<sup>b</sup> |
| $K$ | Carrying capacity, resources | $\mu\text{g chl } a \cdot L^{-1}$ | 25-100<sup>b</sup> |
| $m$ | Loss rate of parasites due to sinking, etc. | $\text{day}^{-1}$ | 4<sup>b</sup> |
| $v$ | Virulence mortality | $\text{day}^{-1}$ | 0.04<sup>a</sup> |
| $w$ | Intrinsic rate of increase, $R$ | $\text{day}^{-1}$ | 0.9<sup>a</sup> |
| $\beta_i$ | Transmission rate, focal hosts $i$ | $L \cdot \text{parasite}^{-1} \cdot \text{day}^{-1}$ | $1.8-5.2 \times 10^{-6}$ <sup>b</sup> |
| $\beta_{av}$ | Average transmission rate, $\sum_{i=1}^n h_i \beta_i$ | $L \cdot \text{parasite}^{-1} \cdot \text{day}^{-1}$ | $1.8-5.2 \times 10^{-6}$ <sup>b</sup> |
| $\sigma$ | Parasite production per infected host | $\text{parasites} \cdot \text{host}^{-1}$ | $1 \times 10^5$ <sup>a</sup> |

<sup>a</sup>Biologically reasonable values (Strauss *et al.* 2015)
<sup>b</sup>Biologically reasonable, see text

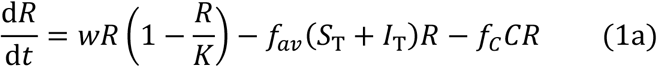

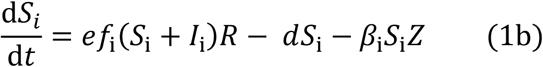

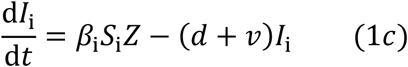

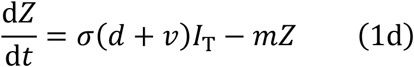

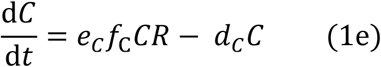

Resources (eq. 1a) grow logistically with intrinsic rate of increase *w* and carrying capacity *K*. They are consumed by the sum of susceptible (*S*_T_) and infected (*I*_T_) hosts of genotypes i = 1 to n 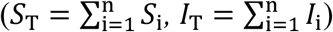. These hosts forage at weighted average rate that depends on the frequency of each genotype 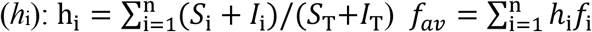. Resources may also be eaten by a competitor species, *C*, with foraging rate *f*_*C*_. Each focal host genotype has fixed trait values, including per-capita transmission rate *β*_i_ (inverse of resistance) and per-capita foraging rate *f*_i_ (linear functional response: Holling 1959; see Fig. A1 for Type II functional response). These traits govern competition for shared resources (eq. 1a) and apparent competition via parasites (eq. 1b).

Both host classes, *S*_i_ and *I*_i_, may convert these resources into susceptible offspring with conversion efficiency *e* (assumed equal among genotypes; eq. 1b). Therefore, transmission is horizontal (not vertical) and infection does not impact fecundity (see Fig. A2 for relaxation of this assumption). Faster foraging genotypes (higher *f*_*i*_) have higher birth rate *ef*_i_*R*. All genotypes experience background mortality rate *d*. Additionally, susceptible hosts can become infected at rate *β*_i_*Z* by encountering parasite propagules while foraging. Foraging and transmission rate positively co-vary, indicating a tradeoff (Fig. 1c). Accordingly, evolving decreased transmission rate (i.e., anti-infection resistance, lower *β*_i_) necessarily lowers foraging rate (*f*_i_) and therefore fecundity (*ef*_i_*R*). While we focus on *f*_i_ (and keep *e* constant), we might have modeled a fecundity-transmission tradeoff of *e*_i_ with *β*_i_, e.g. if high quality offspring require more resources to produce (lower *e*_i_) but have stronger immune systems (lower *β*_i_). In either formulation, the key cost of resistance would be fecundity per resource (*ef*)_i_.

Infection converts susceptible hosts into infected hosts (eq. 1c). These hosts suffer elevated death rate due to virulence of infection, *d*+*v* (where *v* is the added mortality). When infected hosts die, they release *σ* parasite propagules back into the environment (eq. 1d); thus, the parasite is an obligate killer. We do not model parasite evolution as the parasite has not responded to artificial selection (Duffy and Sivars-Becker 2007; Auld et al. 2014). The loss of parasite propagules occurs at background rate *m*. Evolution occurs with changes in clonal frequency solely due to clonal competition (i.e., we assume no mutation, drift, or sexual reproduction).

We also explored the effect of nutrients and a competitor species on host evolution. Nutrients were represented by varying the carrying capacity of the resource (*K*). Competitor density increases as competitors eat resources, with foraging rate *f*_*C*_, convert them into offspring with efficiency *e*_*C*_, and decrease with per-capita death rate *d*_*C*_ (eq. 1e). Because competitors do not directly interact with parasites here, competitors reduce disease via host regulation but not via encounter reduction (Strauss et al. 2016). Competitors were absent in most simulations and analyses (*C* = 0 at *t* = 0), as they were absent in 2/3 of treatments in the motivating experiment.

### Analytical Insight

We gained analytical insight into the evolution of this parasite-host-resource system by deriving an expression for host fitness (decomposed into per capita birth and death rates). Then, we used the continuous time Price equation to determine how transmission rate will evolve (Gandon and Day 2009). Evolution by clonal selection in the presence of parasites depends on the tension between the benefit and cost of resistance. From equations 1b and 1c, one can derive the per-capita rate of change of the total density of a host genotype, i.e., fitness (*r*_i_):

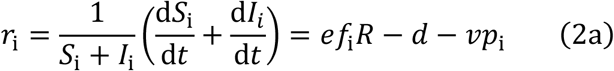

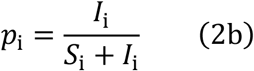

As shown by eq. 2a, fitness is fecundity given conversion efficiency (*e*), foraging rate (*f*_i_), and resources (*R*) minus losses from background mortality (*d*) and infection (*vp*_i_). Prevalence of infection, *p*_i_, is the proportion of hosts of genotype i that are infected (eq. 2b), which is generally higher for genotypes with higher transmission rate (*β*_i_). From this definition of fitness, the rate and direction of resistance evolution can be derived from the continuous time Price equation (as outlined by Gandon and Day 2009):

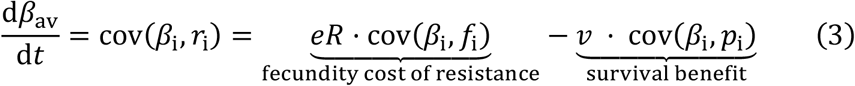

Average transmission rate (*β*_av_; inverse of resistance) changes depending on *β*’s covariance with fitness [cov(*β*_i_,*r*_i_)]; this covariance is simply *β*’s covariance with fecundity minus its covariance with mortality. These terms are closely linked to the fecundity cost and survival benefit of resistance due to the close relationship between covariances and best fit slopes. The slope of fecundity *vs. β*_i_ is the fecundity cost of resistance (lower *β*_i_ corresponds to lower fecundity); *eR*·cov(*β*_i_,*f*_i_) = variance(*β*_i_)·fecundity cost of resistance. Thus, *eR*·cov(*β*_i_,*f*_i_) is the fecundity cost of resistance scaled by variance to determine how the fecundity cost of resistance influences the speed of evolution. Thus, we use *eR*·cov(*β*_i_,*f*_i_) as the scaled fecundity cost of resistance relevant for predicting the speed of host evolution. By the same logic, the second term quantifies the survival benefit of resistance in terms of lower mortality for resistant (low *β*_i_) genotypes, which drives selection for higher resistance (d*β*_av_/d*t* < 0). This formulation clarifies how increased resources can increase the fecundity cost of resistance and drive selection for lower resistance (d*β*_av_/d*t* > 0). Thus, it provides a mechanism by which parasites could increase resources and thus increase selection against resistance.

How much do parasites increase resources? For further analytical insight we use the asymptotic approximation where ecological processes happen much faster than host evolution (i.e., separation of time scales; Jones 1995; Cortez and Ellner 2010). By ignoring feedback from the evolution of transmission rate on host ecology, this approximation focuses on how host ecology at a stable equilibrium influences evolution. We derived this equilibrium for resource density without (*R*^*^_*Z*-_) and with (*R*^*^_*Z*+_) parasites (from eq. 1b and 1c):

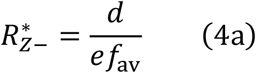

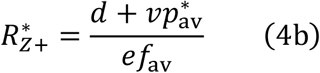

where average prevalence, *p*^*^_av_, is *I*^*^_T_/(*S*^*^_T_ + *I*^*^_T_). Note that these expressions apply in the presence of competitors (who would affect *p*^*\**^_av_) but we focus on interpreting them in the absence of competitors. The symbolic expression for *I*^*^_T_ is too long to easily interpret but putting *R*^*^_*Z*+_ in terms of *p*^*^_av_ proves informative. Combining eqs. 3 and 4 challenges intuitions regarding the effects of parasite virulence or prevalence on parasite-mediated selection. Following conventional theory, one might expect increasing virulence and infection prevalence to strongly select for resistance since it increases the benefit of resistance. Yet, a more virulent and prevalent parasite *could* also increase resources (eq. 4b) and thus the cost of resistance, as outlined above. Indeed, simulations indicate that increased parasite virulence (*v*) increased the benefit [*v*·cov(*β*_i_, *p*_i_)] and cost of resistance (through increased resources). With a steep tradeoff, the cost increased more strongly with *v* (see Fig. A2), selecting for lower resistance and further emphasizing how resource feedbacks can shape patterns of host evolution.

### Simulations

For numerical insight, we parametrized this model with values reasonable for the parasite-host-resource system (see Table 1). We set a poorly characterized parameter (*m*) to make epidemic size biologically reasonable (4 while Civitello et al. 2013 chose 0.5). We used linear tradeoffs with a range of slopes similar to that observed in the experimental system. We used a carrying capacity of the resource (*K* = 100) that was approximately the 95^th^ percentile of algal densities observed in the mesocosm experiment with high nutrients; at low nutrients, we used *K* = 25 (also corresponding to ∼95^th^ percentile algal density for low-nutrient mesocosms). The competitors traits are less well characterized but we chose its death rate to equal that of the focal host (*d*_*C*_ = *d*); because the competitor has a smaller body, we chose it to have a higher conversion efficiency (*e*_*C*_ > *e*) but lower feeding rate (*f*_*C*_ < *f*i for any i) than the focal host. Overall, the model is intended to test key concepts and patterns, not quantitatively fit the experimental epidemics.

We then simulated resource feedbacks and host evolution over the course of epidemics. These simulations used eleven host genotypes. Hosts were evenly distributed with a biologically reasonable range of *β* values (1.78×10^−6^ - 5.16×10^−6^, comparable to trait measurements) and equal starting frequencies (1/11; equal starting variance in *β*_i_ regardless of tradeoff). For simplicity, we used a linear tradeoff with foraging rate as a function of transmission rate. Tradeoff slopes (1,240 parasites·host^-1^ for shallow or 5,944 for steep, see Fig. 1c) were chosen to resemble slopes observed in experimental trait measurements (see below) while intercepts were chosen based on slopes. These choices kept initial, average foraging rate equal regardless of slope [*f*_av_(0)= 0.0344 L·host^-1^·day^-1^]. Hence, steeper slope necessitates lower intercept. Simulations with and without disease were initialized with resources [*R*(0) = 24.2] and hosts [*H*_T_(0) = 59.0] corresponding to their mean densities when parasites were added in the mesocosm experiment. Simulations lasted 125 days after starting without parasites (*Z*-) or with parasites (*Z*+) from a small propagule inoculum [*Z*(0) = 2 × 10^6^ equivalent to spores released from two infected hosts, Figs. 2-3]. We also ran simulations with lower nutrients (by decreasing *K*) and with presence of the competitor (*C*(0) = 7.4 competitors·L^-1^) to recreate those experimental treatments and model their impact on evolutionary outcomes. We averaged the cost of resistance among genotypes over the course of each simulation and computed host evolution from the average transmission rate at the end of simulations compared to the beginning (Δ*β*_av_; e.g., Fig.2).

**Figure 2.**
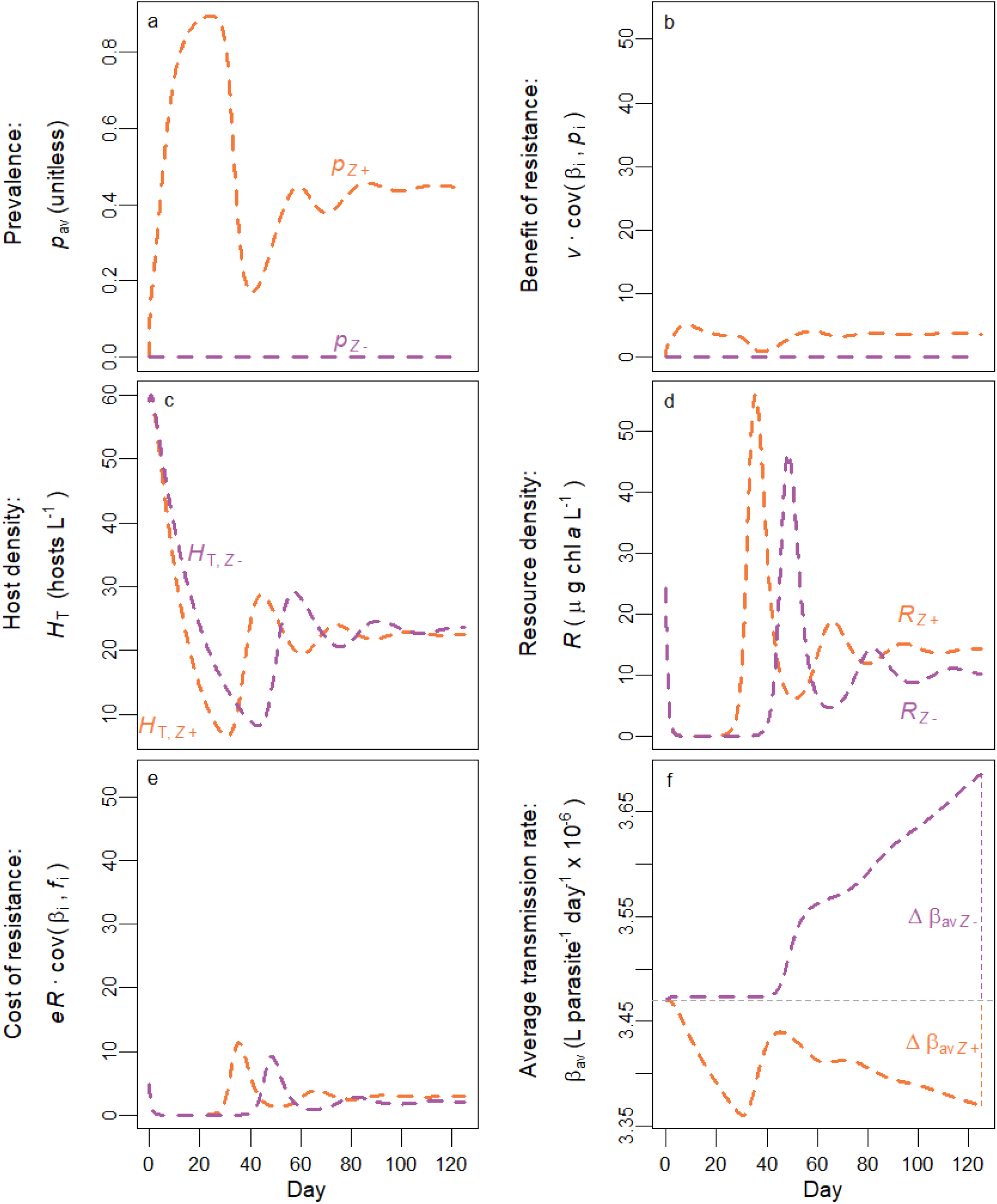
Parasites select for higher resistance when the tradeoff is shallow. (a) Prevalence of infection may be high in the presence of parasites (orange) but is zero in their absence (purple). (b) There is no benefit of resistance in the absence of parasites (purple) but with parasites, resistant genotypes benefit by suffering infection less often [orange; *v*·cov(*β*_i_,*p*_i_) > 0]. Units are L parasite^-1^ day^-2^ × 10^−9^. (c) Parasites also decrease total host density (*H*_T_, orange below purple on average) and (d) increase resource density (*R*; orange above purple on average) in a trophic cascade. (e) Increased *R* makes the cost of resistance higher with disease than without. Units are L parasite^-1^ day^-2^ × 10^−9^. (f) Without parasites, hosts evolve decreasing resistance (*β*_av_ increasing to maximize foraging rate, *f*) relative to their starting resistance (horizontal dashed line). With parasites, hosts evolve increasing resistance. Evolution can be measured as change in *β*_av_ from the beginning to the end of the time series, Δ*β*_av_ (vertical dashed segments). Parameters as in Table 1 (shallow tradeoff slope) with initial conditions of *Z* = 0 or 2 × 10^5^, *R* = 24.2, *S*_*T*_ = 59.0, *I*_*T*_ = 0.

With a shallow tradeoff slope (as in Fig.1c), parasites strengthened selection for resistance in an example simulation. In a population with parasites (*Z*+), prevalence increased rapidly (Fig. 2a), creating a survival benefit of resistance (Fig. 2b) in terms of avoiding infection and corresponding mortality. Parasite-induced mortality decreased host density (orange below purple, on average, in Fig. 2c) and increased resources (orange above purple in Fig. 2d). These increased resources amplified the cost of resistance (orange above purple in Fig. 2e). But this advantage stayed small because of the shallow tradeoff [small cov(*β*_i_,*f*_i_)]. So, parasites increased the survival benefit of resistance (Fig. 2b) more than they increased the fecundity cost (Fig. 2e). Cost and benefit integrated over time determine the change in average transmission rate over the time series 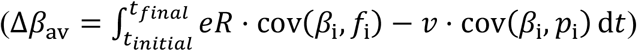. That change in average transmission rate, in turn, provided an overall measure of the direction and magnitude of host evolution. Parasites selected for resistance relative to when parasites were absent (Δ*β* _av, *Z*+_ < Δ*β*_av, *Z*-_). Hence, despite trophic cascades, parasites drove selection toward resistance, at least with a shallow tradeoff.

However, this trophic cascade had a much bigger impact on selection when multiplied by a steep tradeoff (i.e., larger covariance given equal variance in *β*_i_). The patterns for prevalence (Fig. 3a), the benefit of resistance (Fig. 3b), host density (Fig. 3c) and resource density (Fig. 3d) were essentially the same as for the shallow tradeoff. There was a short period of a negative covariance between transmission rate and prevalence; this transient phenomenon arose as highly fecund genotypes made many susceptible offspring that had not yet been infected but were soon to become so. But due to the steeper tradeoff, the cost of resistance (Fig. 3e) was larger and more responsive to resources (compared to Fig. 2e). Specifically, parasites strongly increased the cost of resistance (the product of resources and the tradeoff). These strong costs selected against resistance more strongly in the presence of parasites than in their absence (Δ*β*_av, *Z*+_ > Δ*β*_av, *Z*-_; Fig. 3f) via resource feedbacks.

**Figure 3.**
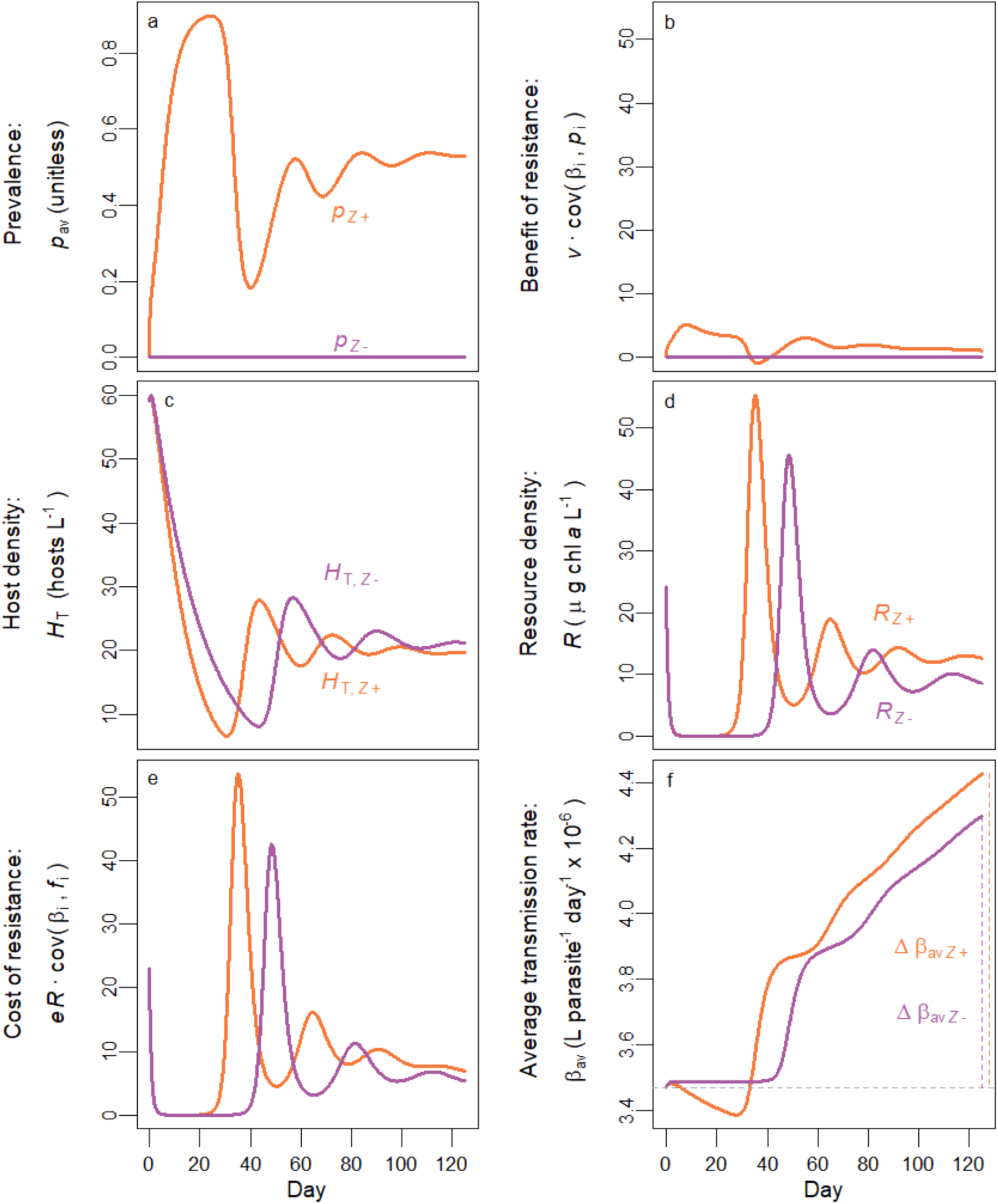
Parasites select against resistance when the tradeoff is steep. (a-d) Follow Fig.2. (e) Increased *R* multiplied by a steep tradeoff makes the cost of resistance much higher with parasites than without (compare to Fig.2e on the same scale). Units are L parasite^-1^ day^-2^ × 10^−9^. (f) Without parasites, hosts evolve decreasing resistance (*β*_av_ increasing to maximize foraging rate, *f*). With parasites, hosts evolve decreasing resistance even faster. Parameters as in Table 1 (steep tradeoff slope) with initial conditions of *Z* = 0 or 2 × 10^5^, *R* = 24.2, *S*_*T*_ = 59.0, *I*_*T*_ = 0.

Ecological factors should shift resource density and the effect of parasites on resources, altering host evolution. At equilibrium, nutrients, represented by the carrying capacity of the resource (*K*), do not increase resource density without parasites (eq. 4a); but with parasites, they increase prevalence (Walsman et al. 2021) so that higher nutrients should increase equilibrium resource density (eq. 4b). Simulations qualitatively agree that nutrients increase resources more when parasites are present (Fig. 4a). The effect of competitors is less straight-forward, but they should decrease resource density. Parasites shift competition between focal hosts and competitors to favor competitors more. Competitors, therefore, should decrease resources more when parasites are present; simulations agree (Fig. 4a and see Fig. A3 for full time series). Thus, ecological factors shape the ability of parasites to increase resources in a trophic cascade.

**Figure 4.**
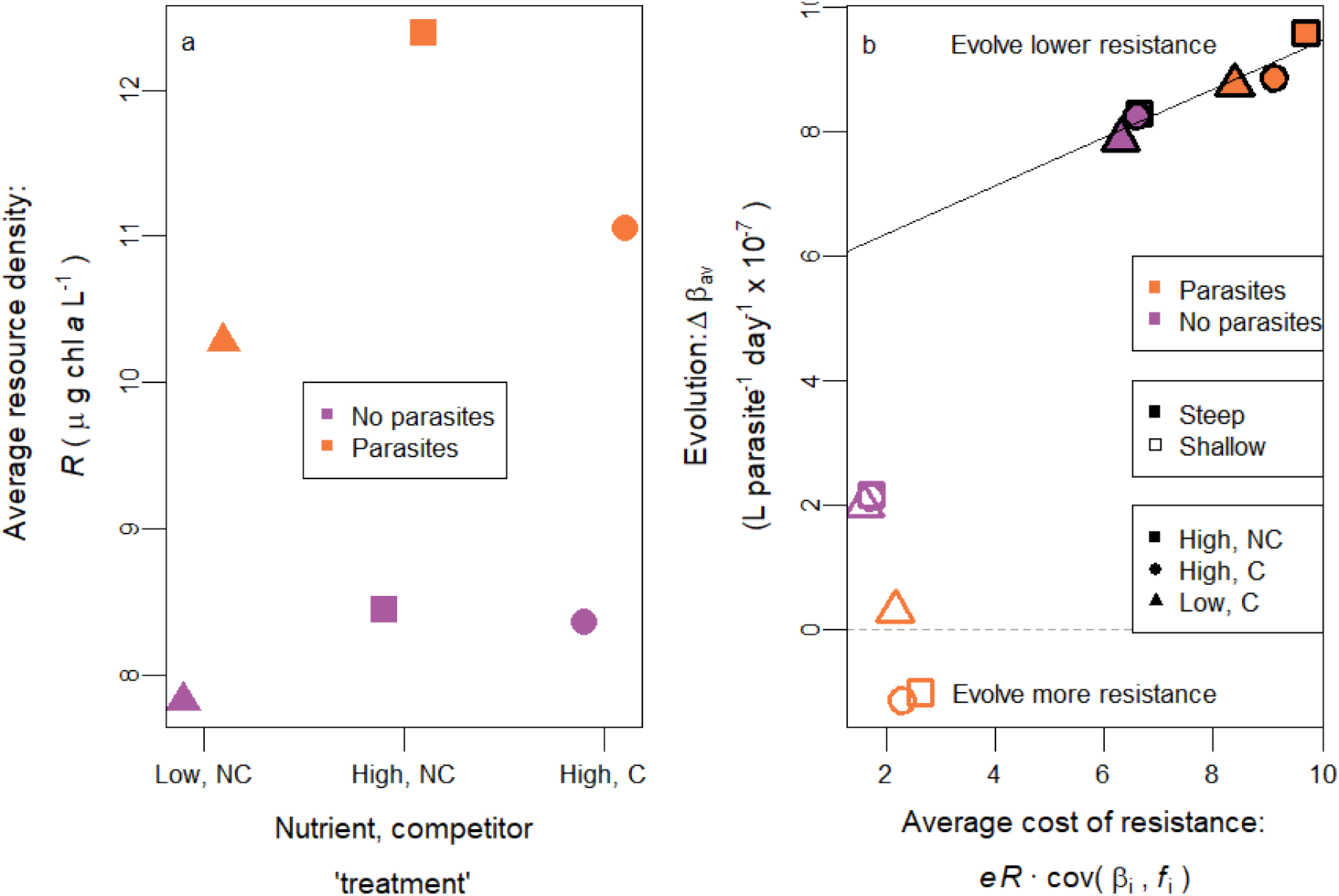
Predictions from simulations of the eco-evolutionary model: parasites can strengthen selection against resistance via increased resources. We simulated the model with high or low nutrients (“High” or “Low”) and competitors present or absence (“C” or “NC”). (a) Simulations show higher resources (*R*, average over time series) with parasites (orange) than without (purple). Low nutrients (triangles) or competitors (circles) decrease *R* and the effect of parasites on *R*. (b) As the cost of resistance (averaged over time series; units are L parasite^-1^ day^-2^ × 10^−9^) increases, host populations with steep tradeoffs evolve toward lower resistance (Δ*β*_av_ > 0; i.e., line has positive slope). If parasites increase the cost of resistance enough, hosts may evolve lower resistance *faster* with parasites than without (e.g., compare orange and purple solid squares). This relationship does not hold for shallow tradeoff populations, which are dominated by the benefit of resistance. Shallow and steep tradeoff slopes correspond to those used in Figs. 2 and 3, respectively. All parameter values as in Table 1.

By decreasing the resource response to parasites, low resources and competitor presence weakened parasites’ ability to increase the cost of resistance, (x-axis, Fig. 4b). Parasites increased the cost of resistance most with high nutrients and competitors absent (horizontal distance between purple and orange is highest for squares in Fig. 4b). The cost of resistance dominated evolution for steep tradeoff populations (solid points) such that higher cost of resistance led consistently to evolution of lower resistance (best fit line shown). For steep tradeoffs, parasites increased the cost of resistance more than they increased the benefit, causing hosts to evolve lower resistance with parasites than without (compare height of solid orange to solid purple, Fig. 4b). This pattern did not hold for shallow tradeoff populations where the costs of resistance were small and did not drive a positive relationship (somewhat weaker, negative relationship). Tradeoff steepness and resource density, together, determine the cost of resistance and how strongly it drives host evolution.

## Eco-evolutionary Experiment

### Experimental Study System

We test our model predictions against experimental populations of dynamically interacting fungal parasites, zooplankton hosts, and algal resources. The focal host, *Daphnia dentifera*, is a dominant zooplankton grazer of algae in North American lakes (Duffy et al. 2004). *Ankistrodesmus falcatus* is a representative, fast-growing alga. The host reproduces clonally, and isoclonal lines vary in transmission rate of the focal parasite and foraging rate (Hall et al. 2010; Hall et al. 2012). Natural host populations often exhibit a strong tradeoff between these two traits (Auld et al. 2013), because hosts incidentally ingest propagules (spores) of the focal parasite, *Metschnikowia bicuspidata* (Ebert 2005). Once consumed, these fungal spores puncture the gut wall of hosts and reproduce in the hemolymph until host death – this obligate killer increases mortality rate. After host death and degradation, parasite propagules are released back into the water column. The parasite has not responded to selection experiments (Duffy and Sivars-Becker 2007; Auld et al. 2014), allowing us to focus solely on evolution of host traits.

### Tradeoff Slopes: Transmission Rates (β_i_) and Foraging Rates (f_i_) of Host Clones

Estimates of transmission and foraging rates of each host clone came from assays on individual host traits. For transmission rate assays, individual hosts of each clonal genotype were previously placed into small (15 mL) tubes and exposed to a range of fungal propagule densities for 8 hours (previously reported by Strauss et al. 2017). Transmission rate was calculated from the probability of infection (see Appendix). For foraging rate assays, individual hosts were placed in 15 mL tubes containing algal resources (*A. falcatus*) for 3-4 hours. We then calculated foraging rate and feeding rate of hosts from the duration and change in algal density (see Appendix). ‘Steep’ and ‘Shallow slope’ tradeoff treatments were created by manipulating which host genotypes were present in experimental populations. Further, we measured foraging rate and feeding rate at two resource densities to test the effect of resources on the advantage of fast foragers (i.e., the fecundity cost of resistance). We fit linear models to determine the steepness of the feeding rate-transmission rate tradeoff among all genotypes at both algal densities to determine if resources increased the cost of resistance.

The tradeoff between foraging rate (*f*_i_) and resistance (inverse of *β*_i_) differed strongly between tradeoff treatments and indicates resources can increase the cost of resistance. The set of genotypes with steep tradeoffs had a slope of 4.17 × 10^3^ parasites·host^-1^ while the weak set had 1.24 × 10^3^ (Fig. 5a). In mesocosms, these tradeoffs differed somewhat depending on empirical genotype frequencies; cov(*β*_i_,*f*_i_) depends on genotype frequencies [mean covariances ± standard error were 37.7 × 10^−10^ ± 3.9 × 10^−10^ L^2^·parasite^-1^·host^-1^·day^-2^ for steep and 1.9 × 10^−10^ ± 2.0 × 10^−10^ for shallow]. Foraging rate determines how quickly a host clears a certain volume of food (*f*_i_ has units of L·host^-1^·day^-1^). The density of food determines how much food is in that volume and thus acquired by a host in a day, i.e., feeding rate (*f*_i_*R* with units of μg chl *a*·host^-1^·day^-1^). If *f*_i_ is constant with respect to food density, then increased *R* will increase *f*_i_*R* and the slope of the *f*_i_*R vs. β*_i_ relationship by whatever factor *R* increases by. We confirmed this by also measuring feeding rate at high algal density (corresponding to roughly 105 *vs*. 10.5 μg chl a L^-1^; Fig. 5b; slope = 4.27 × 10^5^ μg chl *a*·parasite·host^-1^·L^-1^ at high *vs*. slope = 0.48 × 10^5^ at 0.15 low). These feeding rate data indicate that increased resources multiplicatively increase the advantage of low resistance, fast foraging genotypes in the tradeoff (fecundity cost of resistance) as fecundity is related to feeding rate.

**Figure 5.**
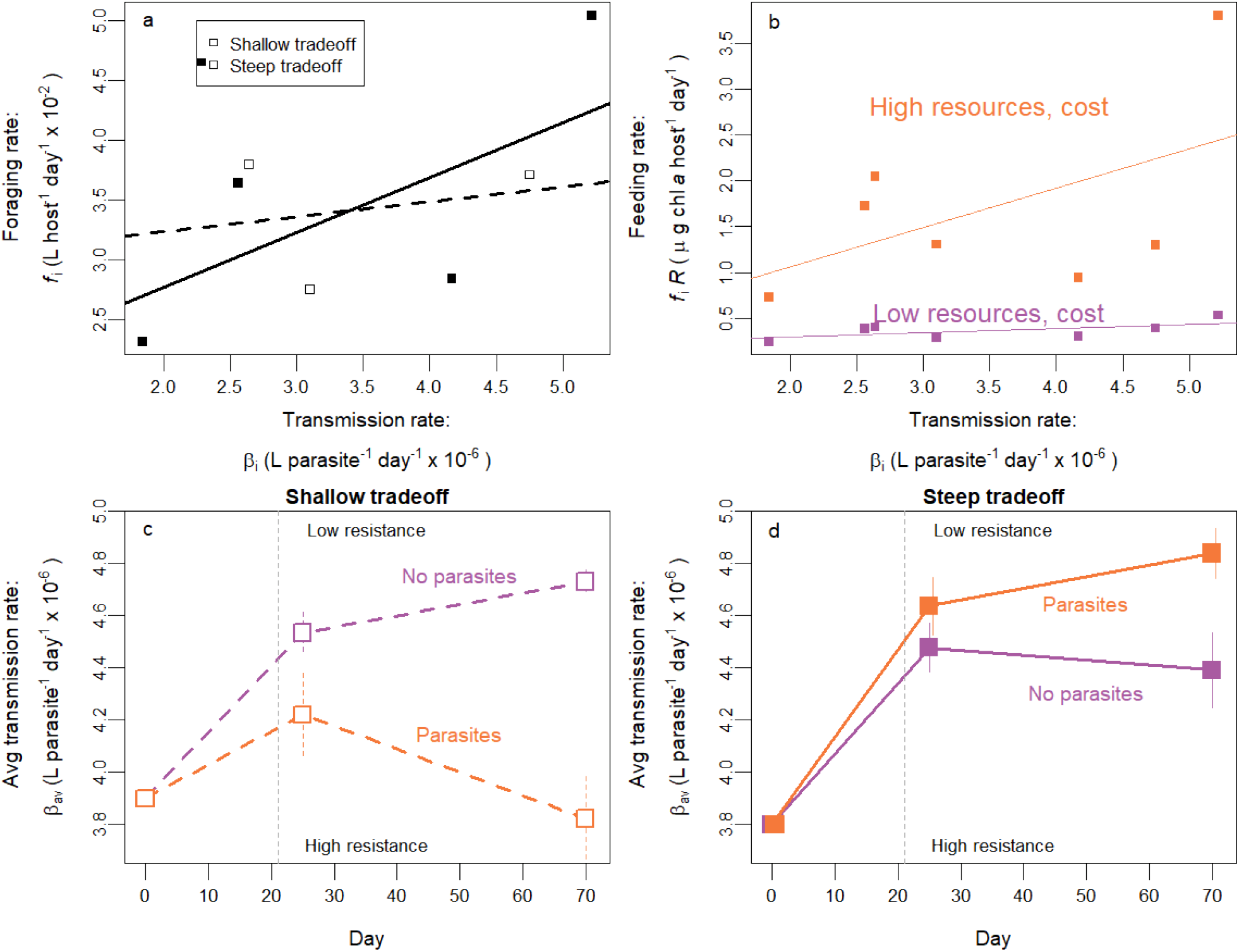
In experimental populations with a steep tradeoff, parasites drove stronger selection for decreased resistance. (a) Clonal host genotypes fall along a tradeoff between transmission rate (*β*_i_, inverse of resistance) and foraging rate (*f*_i_). The slope is steep for the set of seven clones (solid and open points together) but shallow for the subset of three clones (only open points). (b) Measurements at low and very high resource density indicate increased resources increase the slope of feeding rate (*f*_i_*R*), vs. *β*_i_. (c) For a shallow tradeoff, average resistance (inverse of *β*_av_) decreased without parasites (purple line). With parasites, resistance increased during the epidemic (orange line; parasites added at day 21, marked by gray line). Bars denote standard errors. (d) For a steep tradeoff, hosts evolved decreasing resistance but faster with parasites than without. Both c and d show results for all treatments within that tradeoff x parasite combination.

### Other Treatments in the Experimental Communities

Along with tradeoff slope, the mesocosm manipulated nutrients and the presence of a competitor species (Strauss et al. 2017). Host populations were housed in 60 L mesocosms with algal growth stimulated by nutrients and light. Hosts were added at low density on day 0 with all seven genotypes (“strong slope” tradeoff) or a three-genotype subset (“weak slope”). Host populations grew for 21 days before disease-positive (*Z*+) populations were inoculated with 5000 L^-1^ fungal propagules (*Z*). Further, mesocosms’ resources were supplied with low or high nutrients (manipulating carrying capacity of the resource, *K*). Then, at high nutrients only, a competitor species of zooplankton, *Ceriodaphnia* sp, was added. These competitors co-occur in natural lake populations, compete for algal resources, and have low susceptibility to infection by the focal parasite (Strauss et al. 2016).

### Outcomes from Experimental Communities

We determined the evolution of resistance, or lack thereof, from changes in genotype frequencies. Clonal frequency (*h*_i_) was determined for each mesocosm by genotyping up to ten uninfected adult hosts collected from each mesocosm population at days 25 and 70 (Strauss et al. 2017). With these estimates of clonal frequency, we calculated (a weighted average of) transmission rate, *β*_av_ = Σ*h*_i_ *β*_i_, at beginning and end of epidemic (see Appendix for details). We determined the effect of disease on the change in transmission rate using a linear mixed model with change in transmission rate of each genotyped animal as the response variable, parasite presence crossed with time as the fixed effects, and mesocosm ID as a random effect. This formulation focused on change in transmission rate over time and its interaction with parasite presence, accounting for some variation across populations in average transmission rate at the beginning of the epidemic.

Evolution of resistance depended on the interaction of parasites and tradeoff steepness, consistent with our model’s resource feedback mechanism. For a shallow tradeoff, our model predicts hosts should evolve increasing resistance over time with disease and decreasing resistance over time without disease (i.e., an interaction; Figs. 2f, 4b). These trends were observed in mesocosms once epidemics started (gray line in Fig. 5c), although the interaction of parasite presence and time was non-significant (P = 0.610), consistent with the prediction that these effects should be small in magnitude (e.g., note scaling in Fig. 2F). For steep tradeoff populations, host evolved decreasing resistance, and evolved decreasing resistance faster with parasites than without [significant parasite presence x time interaction; P = 0.011; matching Fig. 3f]. Thus, parasites selected against resistance, relative to parasite-free controls, for strong tradeoff populations. Note that genotyping uninfected rather than infected individuals from the experiment, if anything, may have underestimated the magnitude of this effect.

The model predicts that this phenomenon occurred as parasites increased resources and the cost of resistance (eq. 3, Fig. 4b). We highlight the mesocosm results with two example time series. Epidemics (Fig. 6a, orange) decreased host density (Fig. 6b) and increased resource density (Fig. 6c) compared to parasite-free controls (purple). Hosts evolved decreasing resistance without parasites (Fig. 6d; Δ*β*_av, *Z*-_ > 0) and increasing resistance with parasites (Δ*β*_av, *Z*+_ < 0). Steep tradeoff populations had similar patterns (Fig. 6e-g) but increased resources were multiplied by a steeper tradeoff so hosts evolved lower resistance faster with parasites than without (Fig. 6h; Δ*β*_av, *Z*+_ > Δ*β*_av, *Z*-_).

**Figure 6.**
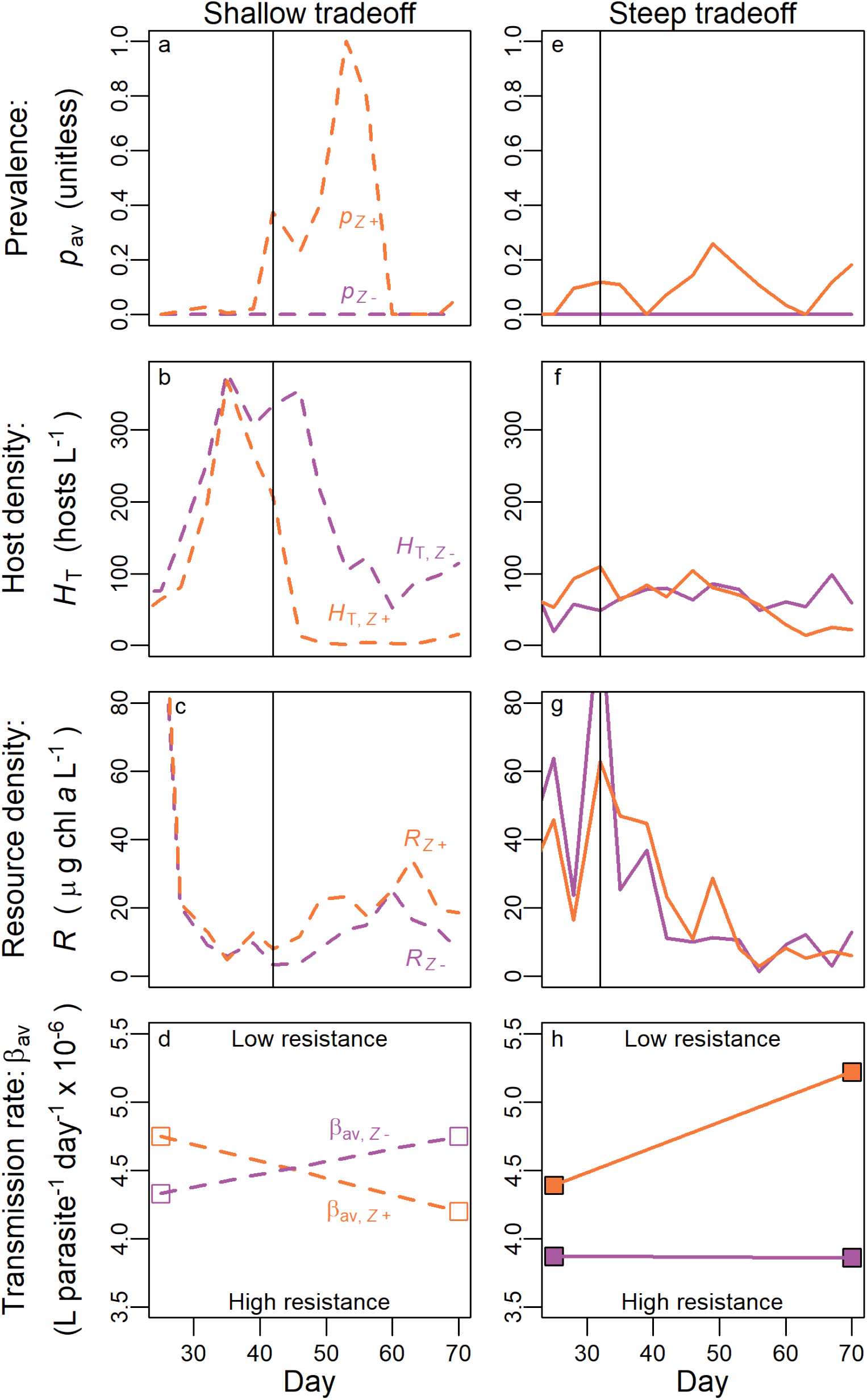
Example experimental populations. One population with parasites (orange, *Z*+) is contrasted to one without (purple, *Z*-) for a shallow tradeoff treatment (dashed) and a steep (solid). (a) Parasites spread after inoculation with a low spore dose on day 21. Population measures span from when parasites first reached a prevalence (*p*) threshold (0.1, here reached at day 42) until the end of the experiment (day 70). Parasites (b) decreased host density (*H*_T_, average = 159.0 for *Z*- and 20.4 for *Z*+) and (c) drove a resource increase (*R*, average = 12.6 for *Z*- and 20.8 for *Z*+). (d) Selection on transmission rate (*β*) is found from changes in *β*_av_ from the beginning (day 25) to the end (day 70) of epidemics (Δ*β*_av, *Z*±_). Hosts evolved increasing resistance (*β*_av_ decreases) with parasites but decreasing resistance without. Results for prevalence (e), host density (f; average = 69.9 for *Z*- and 59.9 for *Z*+), and resource density (g; average = 16.8 for *Z*- and 19.9 for *Z*+) were similar for steep tradeoff populations except increased resources multiplied by the steep tradeoff led to (h) faster evolution of decreasing resistance with parasites than without.

To test model predictions, we also found the average resource density (*R*) for each community. For each disease community, we began integrating resources once prevalence first reached 0.1 (since epidemics take time to affect resource density) until the end of the experiment (day 70). For disease-free populations (*Z*-), we integrated from the mean start date (when prevalence first reached 10%) of epidemics in the corresponding *Z*+ treatment. We used a linear model to determine the effect of nutrient, competitor, and parasite treatment on resource density with parasite x nutrient and parasite x competitor interactions, following model predictions.

Ecological factors affected resource density in the mesocosm communities, as predicted. In the mesocosms, higher nutrients increased resources (P < 0.001; Fig. 7a) while competitors decreased it (P = 0.011). The model predicted parasites would increase resources via trophic cascades (found in a previous, similar experiment; Walsman et al. 2021). The model further predicted a positive interaction with nutrients and negative interaction with competitor presence; these effects trended correctly but not significantly (P = 0.497, 0.284, 0.749, respectively). Thus, the model correctly predicted the trends in average resource densities but not all of these trends were significant.

**Figure 7.**
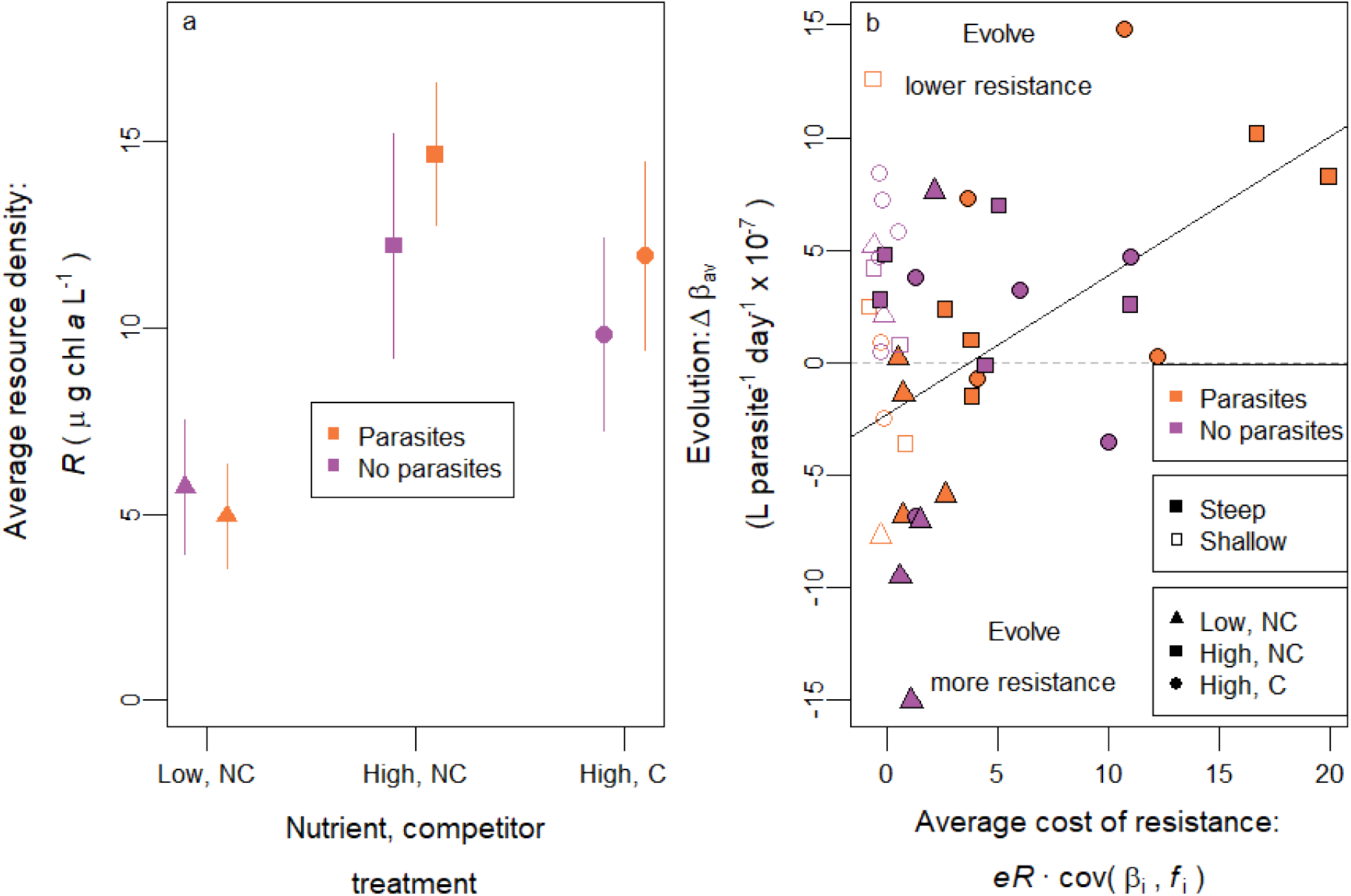
Connection between resources and the costs of resistance indicates parasites selected against resistance through increased resources. (a) High nutrient supply (“HN”) and competitor absence (“NC”) both increased resource density (averaged over time) while parasites did so non-significantly. Bars are standard error. Parasites interacted non-significantly with nutrients and competitors (compare Fig. 4a). Data here is shown pooled for steep and shallow tradeoff populations. Resources were factored into the cost of resistance. (b) As predicted by the model (Fig. 4b), increasing cost of resistance (averaged over time; units are L parasite^-1^ day^-2^ × 10^−9^) selected against resistance (higher Δ*β*_av_) for steep tradeoff populations (solid line). This trend did not hold for shallow tradeoff populations (matching Fig. 4b).

Average resource densities, in turn, were used in calculating the average cost of resistance. We multiplied average resource density by covariance between transmission rate and foraging rate to estimate the average cost of resistance in each host population. We used foraging rate on low resources for this calculation as that resource level was much more representative of average resource density in mesocosm communities (compare Fig. 7a). We regressed the change in average transmission rate on the cost of resistance for all mesocosms with a steep tradeoff treatment with a linear model; we also did the same but for mesocosms with the shallow tradeoff treatment as the model predicted a strong positive response for steep but not shallow. All statistical model assumptions were validated with diagnostic plots (see Appendix and supplementary code).

The model then correctly anticipated that this average cost of resistance, *eR*·cov(*β*_i_,*f*_i_), should correlate positively with resistance evolution, Δ*β*_av_, in populations with steep tradeoffs. The cost of resistance was much higher in steep tradeoff populations (solid points to the right of orange in Fig. 7b) and correlated positively with the increase in transmission rate (P = 0.011; Fig. 7b) as anticipated by the model (Fig. 4b). While most non-disease treatments experienced evolution of lower resistance (due to its fecundity advantages), the largest drops in resistance arose in parasite epidemics. Hence, populations with higher cost of resistance showed large evolution of lower resistance, even (especially) during epidemics. In contrast, the shallow tradeoff populations showed small costs of resistance (i.e., the points were bunched toward the left of Fig. 7b). Not surprisingly, then, no relationship arose between resistance evolution and costs of resistance in those treatments (P = 0.188; compared to Fig. 4b). A strong tradeoff must be multiplied by parasite-driven resource increases for parasites to indirectly select against resistance.

## Discussion

This study showed how parasites can select for or against resistance in their hosts in models and a community-level experiment. In conventional theory, hosts can evolve resistance to avoid infection, which, in turn, can lower mortality (Duffy and Sivars-Becker 2007). During epidemics, then, more resistant strategies attain higher fitness. However, in systems where hosts become infected while foraging, hosts can evolve less resistance when they trade off resistance and foraging/transmission rate (Duffy et al. 2012; Auld et al. 2013; Hall et al. 2010). Our model predicts that this tradeoff drives evolution of decreasing resistance, due to the fecundity cost of resistance, in the absence of parasites (i.e., faster foragers are more fit). But, during epidemics, hosts may evolve increasing or decreasing resistance depending on the fecundity cost and survival benefit of resistance. Critically, the fecundity cost of resistance increases with resource density. Epidemics can increase resource density through trophic cascades. Higher resource density amplifies the advantage of a faster foraging rate, and therefore amplifies the fecundity cost of resistance. However, those same epidemics also increase the survival benefit of resistance (via relatively lower mortality). Consequently, the difference of the cost and benefit then determines evolution of resistance. When a strong trophic cascade in response to parasites accompanies a steep tradeoff, hosts can evolve less resistance with parasites than without.

Generally, most theory anticipates that hosts evolve resistance (Δ*β*_av, *Z*+_ < 0), or at least relatively more resistance (Δ*β*_av, *Z*+_ < Δ*β*av, *Z*-), in the presence of parasites compared to their absence. Without a tradeoff, resistance has no cost. Hence, resistance does not evolve (via selection) without disease (Milks et al. 2002). With disease but still no tradeoff, there is only a survival benefit to resistance. Hence, hosts evolve increasing resistance (model in Duffy and Sivars-Becker 2007). Similar intuitive results arise in models where infection genetics follow ‘matching allele’ mechanisms. In these cases, hosts then evolve resistance to local parasites (Koskella et al. 2011). Evolutionary outcomes become more complex with a fecundity-transmission tradeoff. Parasites may still select for resistance, relative to disease-free conditions, when the system goes to eco-evolutionary equilibrium (Antonovics and Thrall 1994; Best et al. 2017), or if there is no possibility for a resource response (Best et al. 2017). In experiments, similar factors ensure that parasites selects for resistance: there is no tradeoff (Milks et al. 2002), or resource supply is statically provided (e.g., Boots 2011; Duncan et al. 2011). Models and experiments without resource feedbacks should find and have found that parasites select for resistance. Our model and experiment also returned this result, but only when hosts had a weak tradeoff.

In contrast, under certain conditions, epidemics of virulent parasites can actually accelerate the evolution of decreasing resistance. First, the tradeoff mechanism must enable higher resources to increase the fecundity cost of resistance (e.g., Vale et al. 2015), unlike tradeoffs where resources decrease the cost (e.g., Boots 2011). Here, that mechanism hinged on foraging rate since it controlled the exposure component of transmission and the acquisition component of fecundity (Hall et al. 2010; Auld et al. 2013; Hall et al. 2012). Second, resources must respond strongly enough to epidemics. Epidemics can increase resources when heightened mortality of hosts drives trophic cascades (Buck and Ripple 2017). These cascades, in turn, require that hosts control their resources strongly enough to enable resource release and that sufficient time passes for cascades to unfold. We tested these predictions by re-analyzing a mesocosm experiment and calculating the cost of resistance. When clonal assemblages of hosts had strong foraging-transmission tradeoffs, parasites did indeed select for lower resistance (Fig. 5d). Hence, through feedbacks, resources influenced the cost and evolution of host resistance. We found that resources can *increase* the cost of resistance while resources have been found to decrease the cost of resistance in multiple systems. In either case, fluctuations in resources will alter resistance evolution. Given that parasites across systems can drive trophic cascades (Buck and Ripple 2017), the indirect effect of parasites on resources and thus host evolution may have broad importance.

How likely are host-parasite systems to experience trophic cascades that cause parasites to select against resistance? Several factors could influence it. First, many parasites depress host fecundity (examples in Kuris et al. 2008), shrinking the fecundity advantage of less resistant genotypes and strengthening disease-driven selection for resistance (see Appendix). Second, more complex foraging biology (type II or III functional responses: Holling 1959) may increase or diminish opportunities for resources to increase the cost of resistance (see Appendix). Hosts can show power-efficiency tradeoffs such that resource quantity or quality dictate which genotype has the highest fecundity-per resource (Hall et al. 2012). Models could incorporate this additional effect of resources (positive or negative) on the cost of resistance. Third, genotyping enough hosts of known infection status could also determine prevalence for each genotype, enabling estimation of the benefit of resistance (a key component that we could not measure here). Fourth, ecological factors such as competitors may constrain parasite-driven trophic cascades, weaking this mechanism by which parasites select against resistance. Thus, considerations of castration, foraging behavior, the benefit of resistance, and ecological factors for trophic cascades would more broadly determine the likelihood of parasites selecting against resistance.

The likelihood of such a phenomenon would inform how much it should concern disease ecologists. The key ingredients we outlined can guide disease ecologists when looking for parasites selecting against resistance. And that search matters: if hosts evolve less resistance during a large epidemic, parasites may spread more while host density declines lower than otherwise expected. These outcomes could trigger concerns for disease spillover (Power and Mitchell 2004) and persistence of host populations (Ebert et al. 2000). Hence, we need to know if and when parasites and their trophic cascades select against resistance.

## Acknowledgments

O. Schmidt assisted with the trait measurement assays. C. Lively, F. Bashey-Visser, and M. Wade provided valuable feedback on the manuscript.

## Declarations

This work was supported by NSF DEB 1353749 and 1655656 and NSF GRFP awards to J. Walsman and A. Strauss.

## Appendix

### Overview

We cover two topics in this appendix. Section 1 provides further modeling results considering hosts functional response, virulence, and fecundity reduction. These additional results help clarify when parasites should or should not select against resistance. Section 2 further details experimental and statistical methods.

#### Section 1: Further modeling results

##### Resources could increase or decrease the cost of resistance for other host functional responses

Feeding rate may not increase linearly with resource density (Holling 1959). For example, feeding rate may follow a type II functional response = *a*_i_*R*/(*h*_i_+*R*) where *a*_i_ is the maximal foraging rate of a genotype and *h*_i_ is its half-saturation constant. Depending on how these traits covary with resistance (inverse of *β*_i_), increased resources may increase or decrease the cost of resistance. This principle is illustrated most readily with two host genotypes with feeding traits *a*_i_, *h*_i_, and transmission rates *β*_i_. Assume some fixed *β*_1_ > *β*_2_ (and equal, constant conversion efficiencies) so that the fecundity cost of resistance (steepness of *f*_i_-*β*_i_ relationship) is proportional to the difference in feeding rates *a*_1_*R*/(*h*_1_+*R*)- *a*_2_*R*/(*h*_2_+*R*). As a first example, assume the two genotypes have the same maximum feeding rate (*a*_1_ = *a*_2_) but the low resistance genotype’s feeding rate approaches that maximum more quickly (*h*_1_ < *h*_2_). At low resources, the low resistance genotype’s feeding rate will increase rapidly with resources while the high resistance genotype’s feeding rate will increase slowly; thus, increasing resources will increase the cost of resistance (shown as red portion of Fig. A1a). Mathematically, d/d*R* [*a*_1_*R*/(*h*_1_+*R*)- *a*_2_*R*/(*h*_2_+*R*)] decreases from positive to negative as resources increase, given *a*_1_ = *a*_2_ and *h*_1_ < *h*_2_. At high resources, the low resistance genotype’s feeding rate cannot increase further while the high resistance genotype’s feeding rate is still approaching the maximum; thus, increasing resources decrease the cost of resistance (blue portion of Fig. A1a). As a second example, the low resistance genotype has a higher maximum feeding rate (*a*_1_ > *a*_2_) and both genotypes approach their respective maximums with the same half-saturation constant (*h*_1_ = *h*_2_). Now, increasing resources can only increase the difference in feeding rates (all *R* ranges are red in Fig. A1b). Mathematically, d/d*R* [*a*_1_*R*/(*h*_1_+*R*)- *a*_2_*R*/(*h*_2_+*R*)] is always positive, given *a*_1_ > *a*_2_ and *h*_1_ = *h*_2_. These examples demonstrate how non-linear functional forms provide additional complexity for how resources may increase or decrease the cost of resistance.

**Figure A1.**
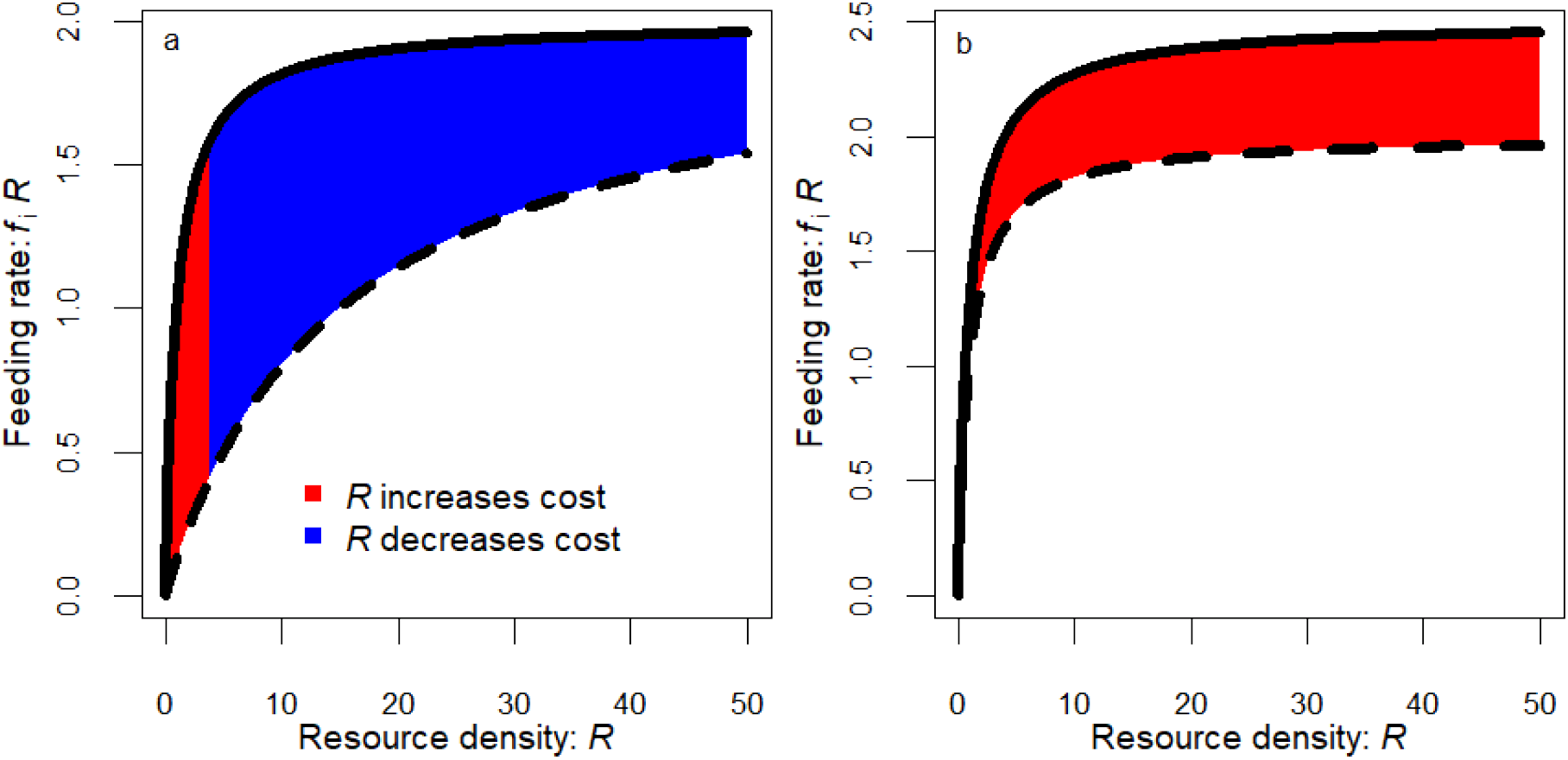
Resources can increase or decrease the cost of resistance with a type II functional response, a_i_R/(h_i_+R). Red shading denotes increasing vertical distance between the two curves as *R* increases. Blue shading denotes decreasing vertical distance between the two curves as *R* increases. In this two-genotype example, the cost of resistance is proportional to the vertical distance between the two curves. (a) The genotype with lower resistance has a feeding rate that saturates faster (solid curve compared to dashed; *h*_1_ = 1 < *h*_2_ = 15) but both genotypes have the same maximum feeding rate (*a*_1_ = *a*_2_ = 2). At low resources, additional resources increase the cost of resistance (red area) as the low resistance genotype’s feeding rate increases more quickly; as the low resistance genotype’s feeding rate saturates, additional resources benefit the high resistance genotype’s feeding rate more than the low resistance genotype’s (blue area). (b) Both genotypes’ feeding rates saturate at the same rate (*h*_1_ = *h*_2_ = 1) but the lower resistance genotype has a higher maximum feeding rate (*a*_1_ = 2.5 > *a*_2_ = 2). Increasing resources only increase the cost of resistance as the low resistance genotype approaches a higher maximum feeding rate than the high resistance genotype.

##### Virulence may decrease or increase resistance

Parasite traits also impact resources, the benefit and cost of resistance, and host evolution. One might expect that increased virulence (here on survival, *v*) should more strongly drive selection for resistance. Higher *v* increases the benefit of resistance [*v*·cov(*β*_i_, *p*_i_)]. This outcome seems especially likely for an obligate-killing parasite (as in eq. 1) because it does not need to keep an infected host alive to transmit. Here, higher virulence does not decrease the probability of infection (unlike for a directly transmitted parasite that needs a living infected host). But as shown in equation 4b (for *R*^*\**^_Z+_), parasites with higher mortality virulence (*v*) could increase resources, depending on how average prevalence (*p*^*^_av_) also responds to higher *v*. Elevated resources can increase the cost of resistance. Thus, more virulent parasites may increase or decrease transmission rate.

Simulating across a range of virulence, our eco-evolutionary model shows that higher virulence increases the cost and benefit of resistance (Fig. A2a). Because we simulated with a steep tradeoff (same as main text), the cost of resistance increased faster with *v* than the benefit; thus, deadlier parasites were more likely to select against resistance (Fig. A2b).

If parasites decrease the fecundity of infected hosts, then the model can be modified slightly (from eq. 1 to eq. A1):

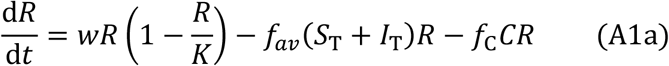

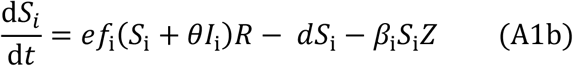

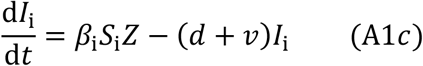

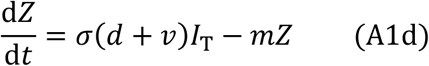

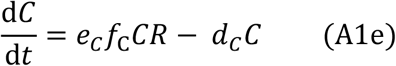

From these equations, we can derive modified versions of fitness, *r*_i_, (compare eqs. 2a) and the dynamics of average transmission rate, d*β*_av_/d*t* (compare eq. 3):

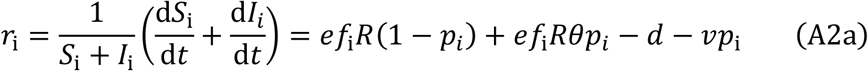

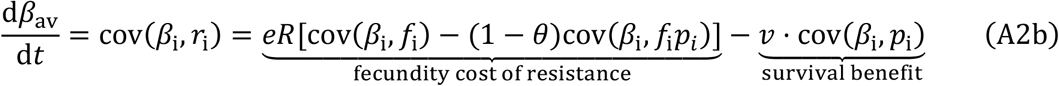

Fitness now depends on contributions of fecundity from uninfected (*ef*_*i*_*R* (1 − *p*_*i*_)) and infected (*ef*_*i*_*Rθp*_*i*_) hosts (eq. A2a). So, fecundity reduction (1-*θ*) decreases the fecundity advantage of low resistance strategies (i.e., “cost of resistance”: first term in eq. A2b) in a way that is linked to the strength of the tradeoff: at low cov(*β*_i_,*f*_i_), *β*_i_ corresponds to higher *p*_i_ but only slightly higher *f*_i_ and thus cov(*β*_i_,*f*_i_*p*_i_) is low while at high cov(*β*_i_,*f*_i_), *β*_i_ corresponds to higher *p*_i_ but and higher *f*_i_ and thus cov(*β*_i_,*f*_i_*p*_i_) is high. Thus, with a steep tradeoff, increasing fecundity reduction (1-θ) strongly reduces the cost of resistance. As a result, higher fecundity reduction enhances the likelihood that parasites will select for resistance (restoring a prediction of more typical eco-evolutionary theory: Fig. A2d).

**Figure A2.**
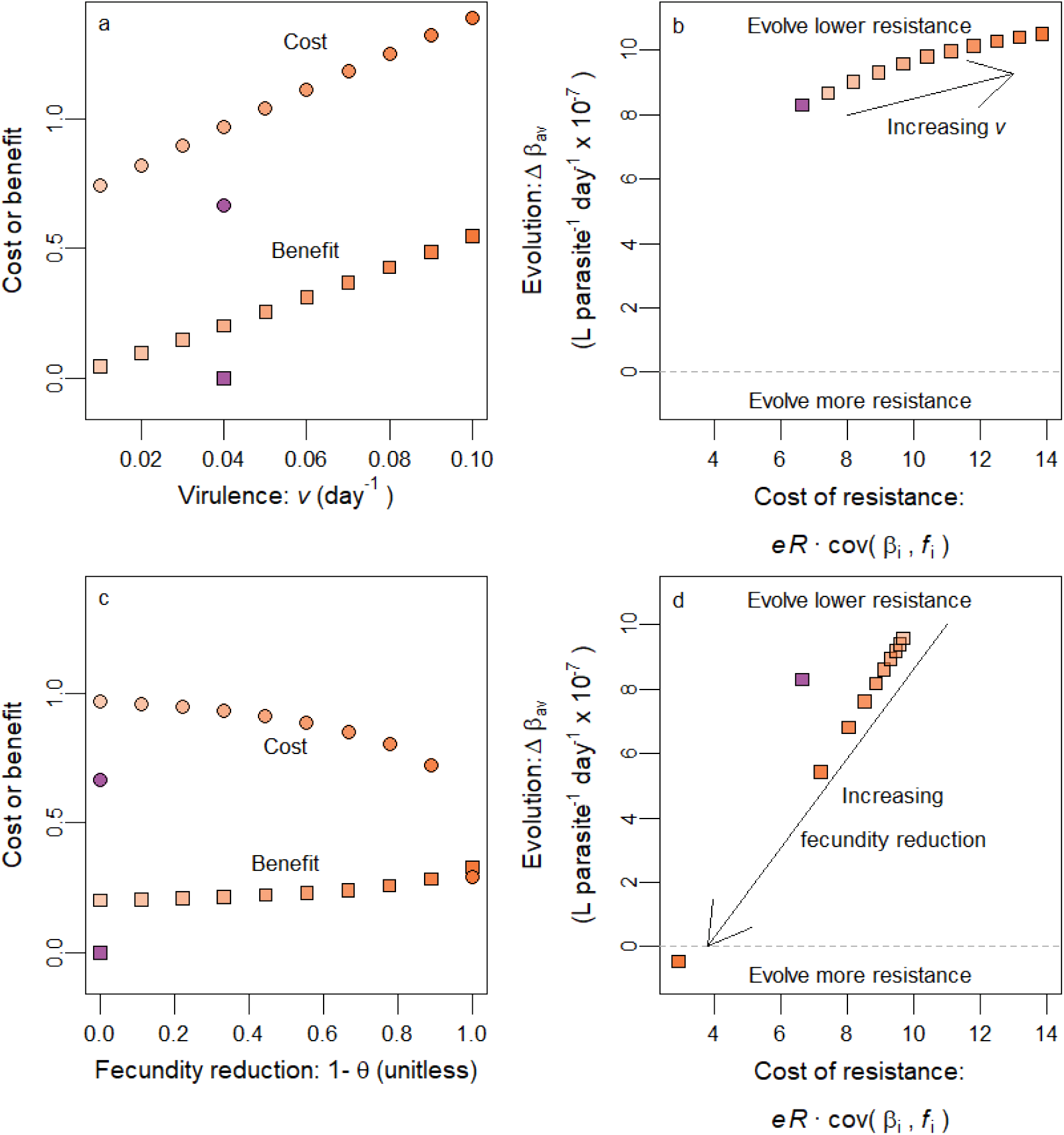
Parasite’s impact on the mortality or fecundity of infected hosts shifts the cost and benefit of resistance, in simulations. (a) As parasites increase in virulence, the benefit of resistance [*v·*cov(*β*_i_, *p*_i_); squares] increases but the costs [*eR*·cov(*β*_i_,*f*_i_); circles] increase faster. Values shown without parasites for reference, here and for other panels (purple; plotted at *v* = 0.04 since that is the default value). More orange shading indicates more harmful parasites. (b) Thus, increasing parasite virulence increases the costs of resistance and drives evolution toward lower resistance (higher *β*_av_). (c) As parasites increase in fecundity reduction (from no castration at *θ* = 0 to full castration at *θ* = 1), the benefit of resistance (squares) increases slightly while the cost of resistance declines rapidly (*eR*[cov(*β*_i_,*f*_i_) – (1-*θ*)cov(*β*_i_,*f*_i_ *p*_i_)] with fecundity reduction; circles). Values shown without parasites for reference (purple; plotted at *θ* = 1 since that is the default value) (d) Thus, increasing fecundity reduction reduces the cost of resistance and drives evolution toward increasing resistance.

#### Section 2: Further empirical detail

##### Measuring transmission and foraging rates of host clones

Transmission rate of individual hosts was measured previously to determine tradeoffs and to infer trait evolution from clonal frequency. Cultures were maintained in high hardness (hh) COMBO (artificial lake water; Baer and Goulden 1998) and ideal conditions (20 deg C, 16:8 light cycle, high food levels [1-2 mg dry mass Anikstrodesmus/L], low density) for three generations (Strauss et al. 2017). This maintenance helped standardize maternal effects. Assays were also performed in hhCOMBO. In brief here: individual hosts were exposed to parasite propagules (either 75, 200 or 393 spores mL^-1^) for 8 hours then transferred to fresh media. Hosts were transferred daily into fresh media until day of death. Dead individuals were examined with a dissecting microscope to diagnose infection. Transmission rate (*β*_i_) was derived from integrating the infection rate [*β*_i_*S*_i_*Z* from eq. 1b: ln(1/(1-*p*_i_)) / (*Z t*_E_) where *p*_i_ is the proportion of hosts that became infected, *Z* is spore dose, and *t*_E_ is time elapsed, c. 8 hours]. Average transmission rate in evolving host populations was determined from clonal frequencies.

We supplemented transmission rate estimates with estimates of foraging rate (*f*_i_) following similar methods. In these assays, we reared cohorts of individuals of a genotype until they reached five days old. Then, we separated them into 15 mL tubes receiving 0.15 or 1.5 mg dry mass/L *Ankistrodesmus falcatus* (∼10.5 or 105 μg chl *a*/L) to determine how feeding rates (closely linked to foraging rates) increased with algal density. Control tubes (five associated with each genotype) were interspersed through the experiment and received the same treatment (i.e., algal density) but without a zooplankton individual. All tubes were inverted approximately every 30 minutes while kept in the dark for 3-4 hours. At the end of the experiment, we removed animals, then measured remaining algae via *in vivo* fluorescence for control and treatment tubes with a Turner Trilogy Laboratory Fluorimeter. With each individual maintained at density *S*, we determined foraging rate *f*_i_ from the algae remaining in the treatment tube (*R*_*f*_) compared to the corresponding control tube (*R*_0_) and the time lapsed, *t*_E_ [i.e. *f* = ln(*R*_0_/*R*_*f*_)/(*S t*_*E*_)]. Feeding rate was computed as feeding rate at the initial algal concentration, *fR*_0_. For a small number of tubes, algal concentration was higher for the treatment tube than the control tube, either due to death of the animal or a molting event (more likely). These treatment tubes were eliminated from the analysis.

##### Population-level experiment and statistical analyses

The methods of the mesocosm experiment have been described in detail previously (Strauss et al. 2017). Genotype frequencies were reported there as well. Hosts were added at approximately equal frequencies on day 0. The starting density of each genotype was the same regardless of tradeoff treatment so the strong tradeoff treatment (21 hosts L^-1^) began with more hosts than the weak tradeoff treatment (8.3 hosts L^-1^). A single genotype of competitors was also added on day 0 for the competitor treatment (2.1 hosts L^-1^). Up to ten animals were sampled from each population (occasionally fewer if necessitated by small populations) at the beginning (day 25) and end (day 70) of experimental epidemics. DNA was extracted from individual animals and five microsatellite loci were amplified by PCR. Microsatellite analysis was performed by sent to the W.M. Keck Center for Comparative and Functional Genomics (University 127 of Illinois at Urbana-Champaign Biotechnology Center, Urbana, IL, USA). Genotypes were identified by comparing their alleles at these loci to known alleles of the isoclonal lines maintained in the laboratory. Some mesocosms (9) were excluded in the final analysis (Fig. 7b) as only one genotype was detected at day 25, precluding a meaningful calculation of cov(*β*_i_,*f*_i_). One more mesocosm was excluded as the prevalence threshold was never reached. This left 30 steep and 20 shallow tradeoff populations.

Not reported previously, were the data and methods for the low nutrient mesocosms. These mesocosms were supplied with 2 μg·L^-1^ phosphorus as K_2_HPO_4_ and 300 μg·L^-1^ nitrogen as NaNO_3_ so that only phosphorus levels differed from high nutrient mesocosms. For low and high nutrient mesocosms, phosphorus and nitrogen were added throughout the experiment to compensate for 5% daily loss from the experimental water column.

We determined the effect of parasites and tradeoff steepness on evolution of transmission rate. Given known transmission rate for each genotype, we used the genotype of each individual as one data point for transmission rate. We regressed transmission rate on time (day 25 or 70) crossed with parasite treatment (present or absent) with each individual mesocosm as a random effect. We did this analysis for the steep tradeoff treatment and the shallow tradeoff treatment. The sign and significance of the interaction describes whether parasites selected for or against resistance (inverse of transmission rate). We conducted these tests using the R package glmmTMB (Brooks et al. 2017). We tested the validity of model assumptions using the DHARMa package (Hartig 2021). While predictions *vs*. residuals plots and tests of dispersion were satisfactory, the data does appear to deviate from a normal distribution. Alternative distributions were tested but were not more satisfactory. All other statistical tests were simple linear models with diagnostic plots checked for the appropriateness of assumptions; plots examined normality, linearity, homoscedasticity, and influential cases. From these diagnostic plots, we determined that these assumptions were met sufficiently.

**Figure A3.**
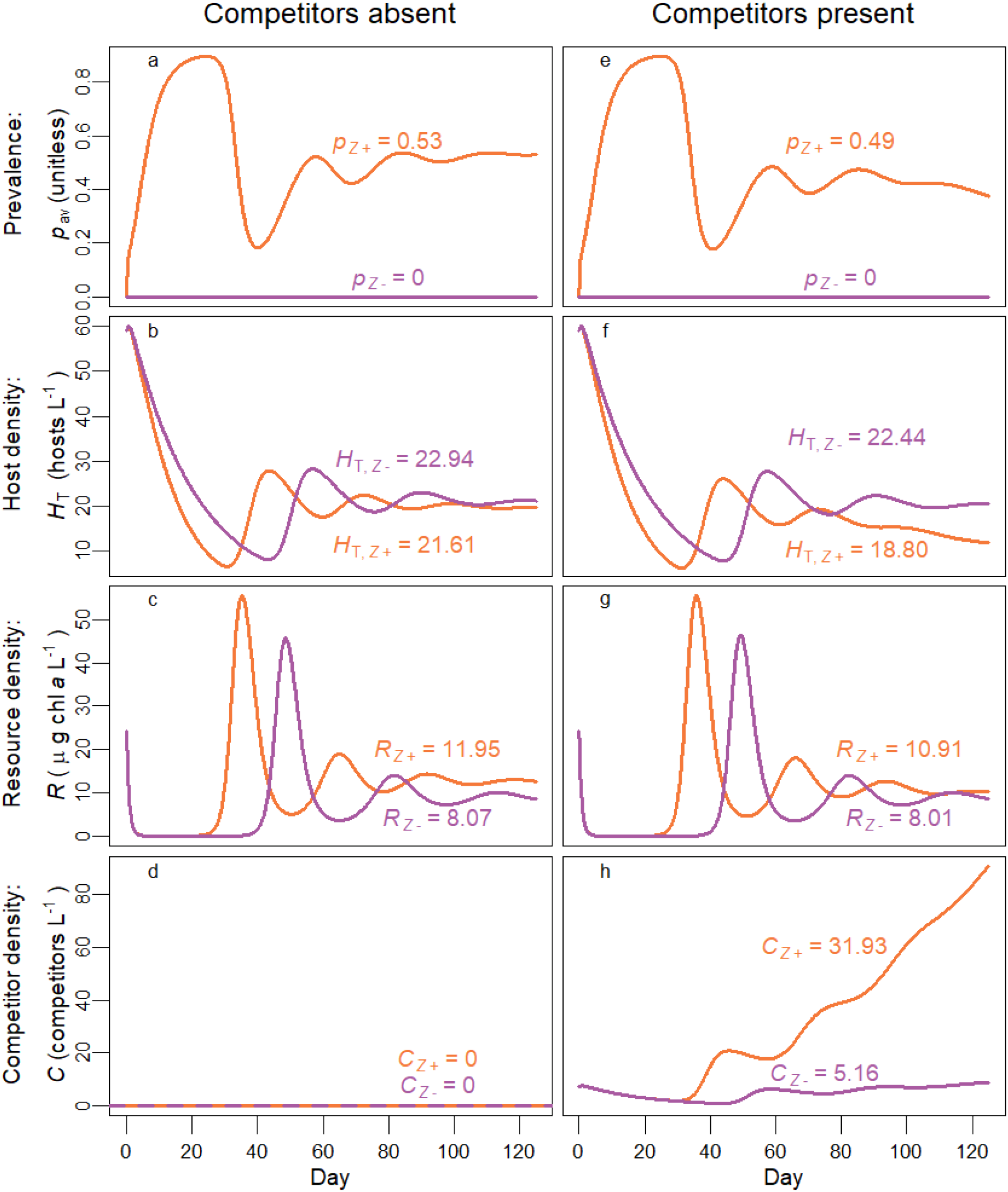
In simulations, competitors decrease host and resource density, especially when parasites are present. Prevalence (a), host density (b), resource density (c), and competitor density (d) are plotted in simulations where competitors are absent (left column; corresponds to simulations shown in Figure 3). When competitors are present (right column), they reduce prevalence (e). They also reduce host density (f), especially in the presence of disease. By grazing, competitors reduce resource density (g), especially in the presence of disease. Competitors attain higher densities (h) when disease is present. Parameter values as in Table 1 with a steep tradeoff.

